# Rare and novel *RELA* variants are common in systemic autoimmunity

**DOI:** 10.1101/2024.11.11.622574

**Authors:** Morgan B. Downes, Sonia B. Nambadan, Joanne Chow, Ainsley Davies, Gemma Hart, Thomas D. Andrews, Nicole Lehmann, Isabella Bales, Alamelu Vengatasalam, Arthur Richard Kitching, Giles Walters, Vicki Athanasopoulos, Simon Jiang

## Abstract

**Objective:** Phenotypic diversity of autoimmune diseases presents an ongoing diagnostic and drug development challenge for clinicians and scientists. Recent discovery of mutations in *RELA* (encoding RELA) in patients with different diagnoses has highlighted that different pathogenic molecular mechanisms are at play and may explain the observed phenotypic diversity. We identified seven novel/rare *RELA* variants in patients with autoimmune diseases and examine the functional consequences on immune signalling.

**Methods:** Wild type and mutant RELA proteins were ectopically expressed in HEK293 cells. Western blot and NF-κB/IFNβ luciferase reporter assays were used to determine RELA expression and transcriptional activity, respectively. In patients (n=3), B and T cell populations were examined via flow cytometry and NF-κB and interferon stimulated genes in PBMCs were assessed via qPCR following toll-like receptor activation.

**Results:** RELA^I250V^, RELA^R295H^ and RELA^E3*^ displayed a loss in NF-κB transcriptional activity. Comparative to RELA^WT^, RELA^I250V^ protein expression was reduced. Two variants, *RELA*^*I250V*^ and *RELA*^*R295H*^, induced hyperactivation of the *IFNβ* promoter. An elevated IFN gene signature was not detected in patient PBMCs following toll-like receptor activation, however the patient heterozygous for I250V had elevated *IFNβ* transcripts at baseline and after TLR7/8 activation. A reduction in transitional, unswitched memory and memory B cell and cTfh (CCR6-CXCR3-) T cell subsets was shared by the patient group.

**Conclusion:** We expand upon the clinical syndromes linked to RELA dysfunction and uncover rare and novel variants that have distinct functional effects on gene transcription downstream of *NF-κB* and *IFNβ* promoter elements. These findings reinforce an important role for RELA in a range of autoimmune and autoinflammatory diseases.

## Introduction

Autoimmune diseases encompass a diverse group of conditions affecting approximately 10% of the population worldwide. The clinical manifestations of disease are highly variable which reflects underlying pathophysiologic heterogeneity. While this heterogeneity significantly impairs diagnosis and treatmen^1^, identifying the genetic drivers of disease and subsequent mechanisms has the potential to improve treatment outcomes.

Nuclear factor κB (NF-κB) is a family of transcription factors that control innate and adaptive immune processes via their regulation of >500 genes. Canonical and non-canonical NF-κB gene targets are involved in cell development, proliferation, inflammation, survival and adaptive immune cell function ^2^. Mutations in NF-κB factors, signalling partners and regulators have been described in patients with immunodeficiency with concomitant autoimmune and/or autoinflammatory manifestations^3^. It has been recently reported that variants in NF-κB factor *RELA* further link dysregulated NF-κB signalling and autoimmunity^4-6^. *RELA* encodes the Rel protein p65/RELA which can homodimerize or form a heterodimer with p50. The RELA/p50 heterodimer remains inactive, sequestered in the cytoplasm until stimulation-dependent post-translational modification signals allow RELA/p50 to translocate into the nucleus and initiate target gene transcription^7^. The recent discovery of *RELA* variants in patients with autoimmune diseases has not only extended the clinical phenotypes assigned to *RELA* variants but also highlighted mechanistic differences that contribute to RELA dysfunction ^6^. It is important, therefore, that *RELA* variants be comprehensively characterised in patients with autoimmune diseases, as this knowledge will have valuable application within clinical settings to improve patient care. Here, we aimed to identify and characterise the contribution of pathogenic *RELA* variants in various autoimmune conditions.

## Methods

### Participants

Patients were recruited as part of the Centre for Personalised Immunology (CPI) program. The study was approved by Australian Capital Territory Health and Australian National University Human Research Ethics Committees. All participants provided written informed consent in accordance with the Declaration of Helsinki.

### Human peripheral blood mononuclear cells (PBMCs) preparation

PBMCs were isolated by density gradient centrifugation using Lymphoprep™ separation medium (StemCell Technologies) and cryopreserved in fetal bovine serum (FBS, Gibco) with 10% dimethyl sulfoxide (DMSO, Sigma).

### Whole-exome sequencing (WES) and variant scoring

DNA samples were enriched using the Human SureSelect XT2 All Exon V4 Kit and sequenced using the Illumina HiSeq 2000 system. Bioinformatics analysis was performed at the John Curtin School of Medical Research (JCSMR), ANU. Raw sequence reads were aligned to the human reference genome (GrCh37/38) and single-nucleotide variants and small insertions and deletions called using the Genome Analysis Toolkit. *RELA* variants were identified via a search for ‘de novo’, coding, novel or rare (MAF <0.001) variants among patient sequence data. PolyPhen-2, SIFT, and CADD were used to predict functional effects.

### Dual luciferase assays

HEK293 cells were transfected with the indicated plasmids, 45 ng pNiFty (lnvivoGen) or 45ng IFNβ (a gift from J. P.-Y. Ting, University of North Carolina) firefly luciferase plasmids and 5ng pRL-CMV *Renilla* control plasmid (Promega). 24 hours post-transfection the dual-luciferase assay was performed using Luc-Pair Duo-Luciferase HS Assay Kit (GeneCopoeia). Luminescence was measured using the LUMIstar Omega plate reader (BMG Labtech).

### Transfection, cell culture and western blot

HEK293 cells (ATCC #CRL-1573, Invitro Technologies) were transfected with the relevant plasmids using Lipofectamine 2000 (Thermo Fisher Scientific) following the manufacturer’s protocol. For stability tests, transfected cells were treated with 20 µM MG-132 (Sigma) for 6 hours. Whole-cell extracts (WCE) were prepared using RIPA lysis buffer containing PhosSTOP (Roche) and cOmplete protease inhibitor (Roche). Proteins were separated via denaturing SDS-polyacrylamide gel electrophoresis and transferred to nitrocellulose membranes. Membranes were probed with the relevant primary and secondary antibodies, developed with Clarity Western ECL Substrate (Bio-Rad Laboratories) and imaged on ChemiDoc (Bio-Rad Laboratories). Quantification of band densities was performed using ImageLab software (version 6.1.0, Bio-Rad Laboratories).

### Flow cytometry

2×10^6^ PBMCs per sample were stained with antibodies against the relevant proteins, and with ViaDye Red (Cytek R0-00008), and fixed with 1% paraformaldehyde (Thermo Fisher Scientific) in PBS. Data was collected on a three-laser Cytek Northern Lights (Cytek Biosciences, Inc.).

### Statistical analysis

Statistical analyses were performed using GraphPad Prism 10.3.0 (GraphPad Software). Statistically significant differences are indicated as p≤ 0.05. Mann–Whitney test and one-way ANOVA with Dunnett’s post hoc analysis were used to analyze immunophenotyping results and luciferase data, respectively.

Further details are provided in the Supplementary Methods

## Results

### Identification of rare and novel *RELA* variants

To identify potential pathogenic *RELA* variants we investigated a cohort of (n=1097) patients with a range of autoimmune diseases including systemic lupus erythematosus (SLE), rheumatoid arthritis (RA), and antineutrophil cytoplasmic antibody (ANCA) vasculitis, who had undergone whole exome sequencing and variant analysis as described previously^8^ As rare (minor allele frequency (MAF)<0.005) variants are known to be enriched for significant phenotypic effects^9^, we identified all rare and novel missense *RELA* variants. We identified n=7 rare and novel heterozygous missense variants in *RELA* (Supplementary Table 1) in seven patients with autoimmune diseases (Figure 1A, Supplementary Table 1). Variants were confirmed by Sanger sequencing for patients where PBMC samples were available (Figure 1B).

**Figure 1.**
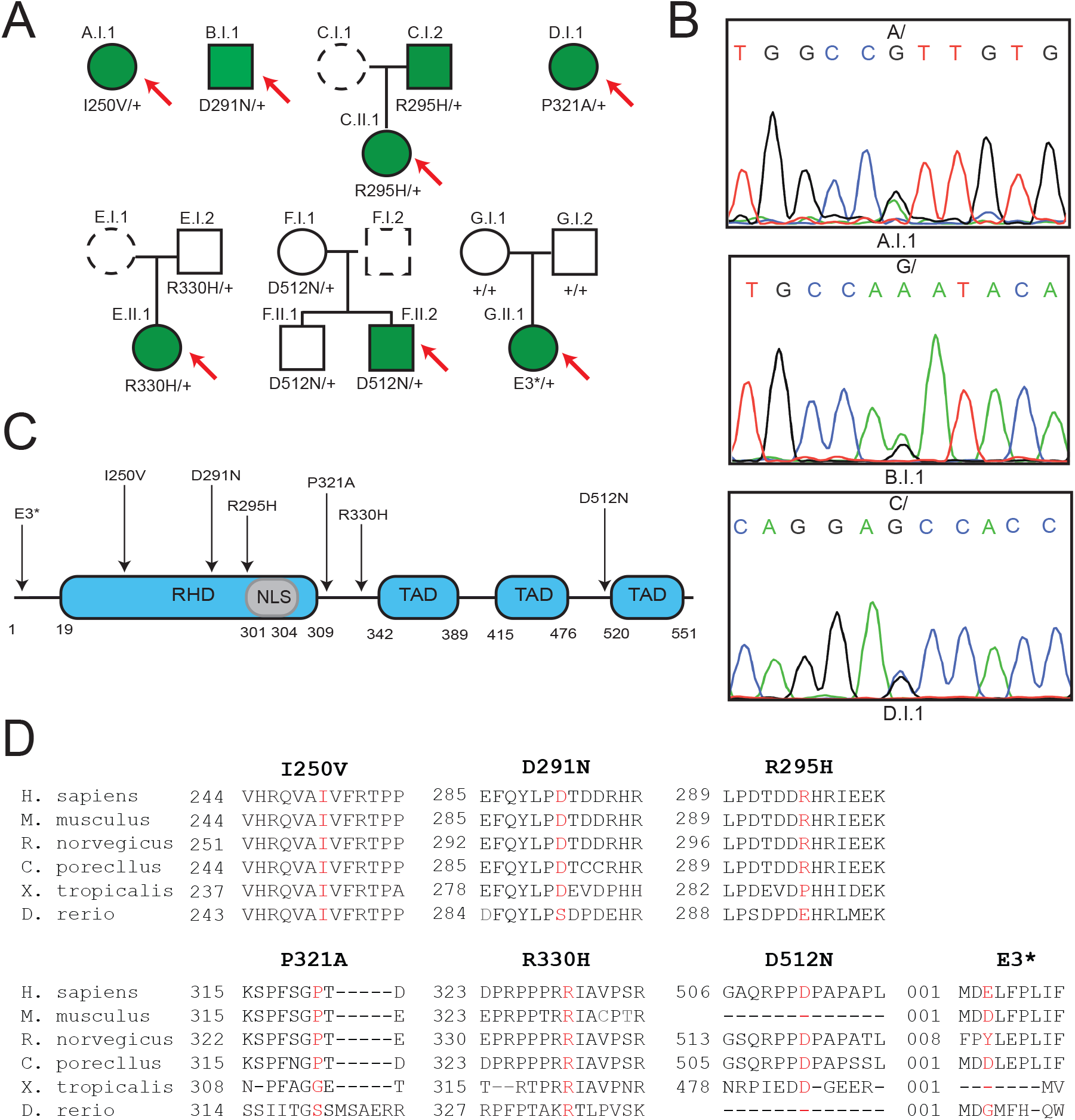
Novel and rare *RELA* variants identified in patients with autoimmune diseases. (A) Pedigrees. Affected (filled symbols), unaffected (unfilled symbols), proband (red arrow), dashed symbols (not genotyped). (B) Sanger sequencing chromatograms. (C) Schematic of RELA protein. Locations of mutated residues indicated by arrows. RHD - Rel Homology Domain, NLS - Nuclear Localisation Sequence, TAD - Transcriptional Activation Domain. (D) Conservation of mutated residues across species.

We identified two novel variants encoding a missense change of isoleucine to valine at amino acid position 250 (*RELA*^*I250V*^) and glutamic acid to a premature stop codon at residue 3 (*RELA*^*E3\**^). The remaining patients carried rare missense variants encoding changes to residues 291 aspartic acid to asparagine (*RELA*^*D291N*^, MAF:0.004278), 295 arginine to histidine (*RELA*^*R295H*^, MAF:0.000261), 321 proline to alanine (*RELA*^*P321A*^, MAF:0.000001239), 330 arginine to histidine (*RELA*^*R330H*^, MAF:0.000007435) and 512 aspartic acid to asparagine (*RELA*^*D512N*^, MAF:0.000004997).

The rare *RELA* SNVs did not segregate to any particular protein domain and were found across the length of the RELA protein; *RELA*^*I250V*^, *RELA*^*D291N*^ and *RELA*^*R295H*^ reside within the Rel Homology Domain, whilst *RELA*^*P321A*^, *RELA*^*R330H*^ and *RELA*^*D512N*^ lie in the disordered C-terminal regions (Figure 1C). Each missense variant alters a highly conserved residue (Figure 1D) and are predicted *in silico* to be deleterious or possibly damaging (Supplementary Table 2). Supplementary Table 3 includes measurements of structural features of each identified amino acid substitution. The I250V substitution is predicted to be mildly destabilising in a buried and ordered part of the structure. Potentially for a protein of RELA’s length, this may suggest a functionally important effect. For the other substitutions, found in intrinsically disordered or surface exposed regions of the protein, the substitutions are not predicted to have stability effects or large changes of physiochemical properties. In summary, we identified four rare or novel missense variants predicted to be deleterious in patients with autoimmunity.

### Autoimmune-associated *RELA* variants are loss of function

We next sought to test the impact of these missense SNVs on RELA function. We overexpressed WT and mutant RELA cDNA plasmids in HEK293 cells and measured their transcriptional capacity by *NF-κB* luciferase reporter assay. For overexpression assays the cDNA plasmid utilized coded for the 448 amino acid RELA isoform (ENST00000612991.4), therefore the D512N variant is herein denoted as D409N and relates to the same residue. As expected, the RELA^E3*^ resulted in complete loss-of-function (LoF), and RELA^I250V^ and RELA^R295H^ proteins showed significantly reduced NF-κB transcriptional activity (Figure 2A). The most abundant canonical NF-κB dimer is the RELA-p50 heterodimer^10^. Although HEK293 cells express endogenous p50, we co-transfected p50 so that ectopic levels were similar to overexpressed RELA to try to maintain stoichiometric ratios and analyse heterodimer activity more accurately. We identified RELA^I250V^ and RELA^R295H^ as hypomorphs, with reduced activity as heterodimers (Figure 2A). Together these data identify three LoF *RELA* alleles in three patients with autoimmunity.

**Figure 2.**
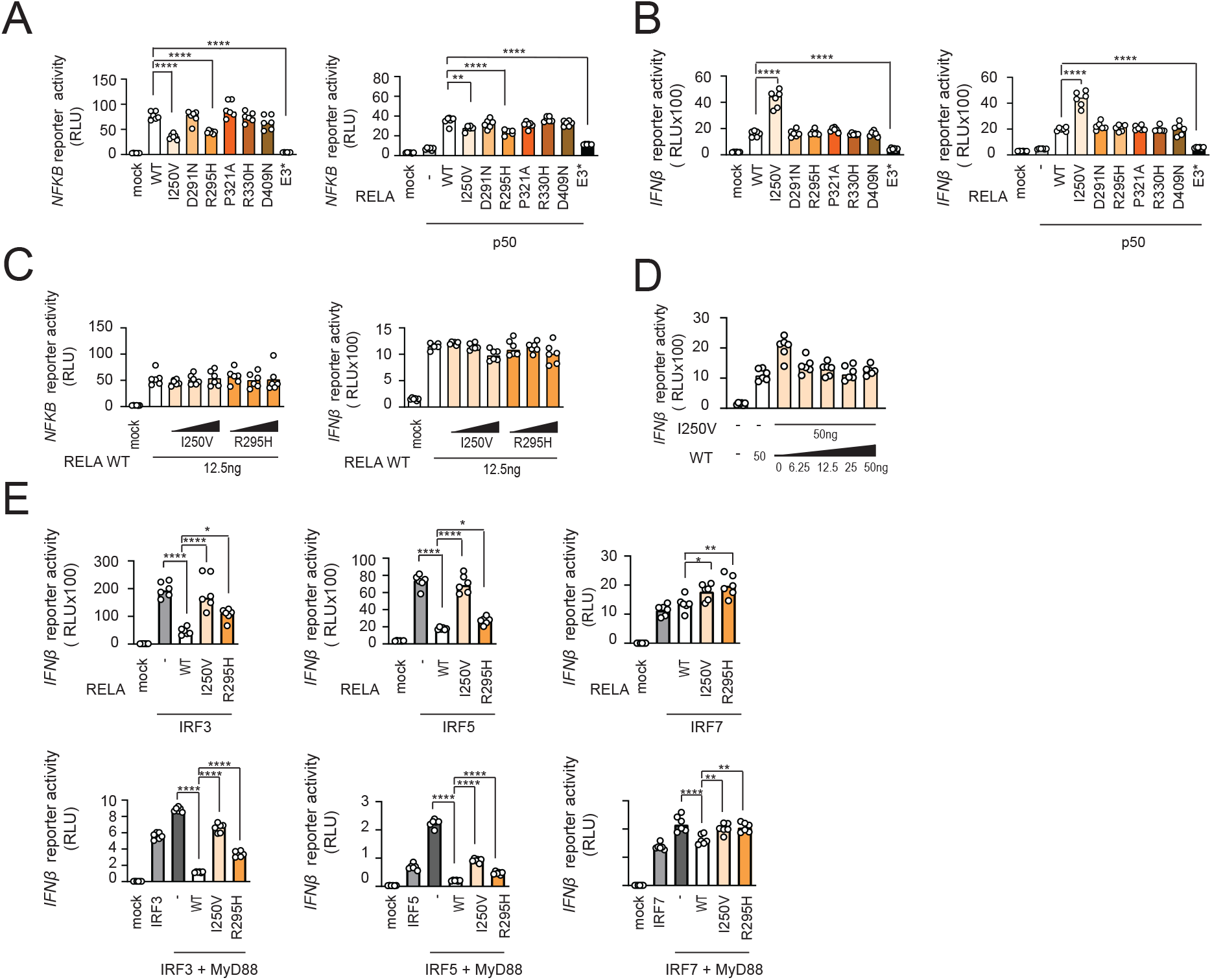
Loss of function *RELA* alleles permit increased IFNβ promoter activity. (A) NF-κB luciferase reporter activity in HEK293 cells transfected with 25 ng of *WT* or *MT RELA* plasmids alone or 12.5ng *WT* or *MT* RELA plasmids with 50 ng *p50*. Graphs represent 2-3 independent experiments. (B) IFNβ luciferase reporter activity in HEK293 cells transfected with 50 ng of *WT* or *MT RELA* plasmids alone or with 50 ng *p50* plasmid. Graphs represent 3-4 independent experiments. (C) NF-κB and IFNβ luciferase reporter activity in HEK293 cells transfected with 12.5 ng *WT RELA* and increasing amounts of *MT RELA* (12.5, 25, 37.5 ng). Graphs represent 2-3 independent experiments. (D) IFNβ luciferase reporter activity in HEK293 cells transfected with 50 ng *RELA*^*I250V*^ and increasing amounts of *RELA*^*WT*^. (E) IFNβ luciferase reporter activity in HEK293 cells transfected with 50 ng of *RELA WT* or *MT* and 50 ng of either *IRF3, IRF5* or *IRF7* plasmids in the absence or presence of 25 ng *MyD88* plasmid. Graphs represent 2-3 independent experiments. All graphs display median values, RLU - relative light units.

Truncating *RELA* variants are reported in patients with an enhanced type I IFN signature ^6^ and RELA is a known co-regulator of type I IFN production forming part of the IFN enhanceosome, an assembly of factors which recruit coactivators and chromatin remodelling proteins to promote *IFNβ* promoter activity ^11^. Patients A.I.1 and E.II.1 were diagnosed with SLE, a prototypical type I interferon-mediated disease ^12^ (Supplementary Table 1), and we therefore examined the impact of *RELA* variants on *IFNβ* promoter activity via luciferase reporter assays. In HEK293 cells overexpressing RELA alone, RELA^I250V^ displayed enhanced *IFNβ* promoter-driven luciferase activity (Figure 2B). Elevated activity by RELA^I250V^ was also evident when p50 (Figure 2B) and the IFNβ enhanceosome coactivator p300 were co-expressed (Supplementary Figure 1A). Wild type levels of transcriptional activity were observed in cells transfected with *RELA*^*R295H*^ alone, however this was elevated in the presence of p300 (Supplementary Figure 1A). Thus, RELA^I250V^ and RELA^R295H^ behave as neomorphs, losing transcriptional activity on cognate *NF-κB* sites and promoting activity on the *IFNβ* promoter.

Recently reported heterozygous LoF *RELA* variants associated with a type I interferon signature in patient blood were found to exert a dominant-negative effect on NF-κB transcriptional activity ^6^. All our patients are heterozygous for the identified *RELA* variants. We therefore tested for a potential dominant-negative effect of RELA^I250V^ and RELA^R295H^ proteins on WT RELA. Increasing quantities of LoF RELA did not suppress the transcriptional activity of RELA^WT^ protein on the NF-κB promoter (Figure 2C). Similarly, the LoF RELA alleles did not exert a dominant-negative effect on *IFNβ* promoter activity (Figure 2C). Of note, the elevated activity of RELA^I250V^ on the *IFNβ* promoter was no longer evident in the presence of RELA^WT^ at all tested doses (Figure 2D). RELA^WT^ complementation rescued the dysregulated IFN pathway activation conferred by RELA^I250V^ at a ratio as small as 1:8. This effect may be due to differential DNA binding affinities between RELA WT and MT proteins or it suggests that RELA plays a regulatory role at the site of IFN transcription. Together this indicates that *RELA*^*I250V*^ and *RELA*^*R295H*^ are non-dominant LoF alleles affecting NF-κB and IFN signalling.

### Neomorphic *RELA* variants alter IRF-dependent activation of IFN signalling

Interferon regulatory factors (IRFs) are transcription factors that are strong drivers of IFN signalling ^3^. Given that two LoF *RELA* alleles enhanced interferon promoter activity we assessed their impact on interferon regulatory factor 3 (IRF3), interferon regulatory factor 5 (IRF5) and interferon regulatory factor 7 (IRF7)-dependent *IFNβ* promoter activation. *RELA* plasmids were co-transfected with *IRFs* alone and in the presence of *MyD88*, an adaptor molecular that facilitates IRF activation. RELA^WT^ suppressed *IFNβ* reporter activity in the presence of *IRF3* and *IRF5* (Figure 2E), but not in the presence of *IRF7* alone. RELA^WT^ repressed *IFNβ* in cells expressing IRF7 and MyD88 (Figure 2E), but to a less extent than for *IRF3* and *IRF5* in the presence of MyD88. Interestingly, the suppressive action of RELA on IRF3, IRF5 and IRF7 activity was abolished by the LoF I250V variant and partially lost with *RELA*^*R295H*^. In the presence of IRF7 alone, RELA^I250V^ and RELA^R295H^ increased *IFNβ* reporter activity. RELA contains IRF binding motifs which allow direct interaction with IRF3, 5 and 7 ^13^. However, the RELA isoform used in luciferase experiments lacks the IRF binding motifs, as demonstrated by the failure to detect IRF7 interaction following RELA immunoprecipitation (Supplementary Figure 1B). Therefore, the suppressive action of RELA on IRF activity is likely due to an indirect mechanism. In summary, RELA^I250V^ and RELA^R295H^ proteins lose the ability to negatively regulate IRF activity on the *IFNβ* promoter.

### Altered stability of mutant RELA proteins

We hypothesised that RELA LoF could result from altered protein levels. We overexpressed wild type and mutant RELA in HEK293 cells and demonstrated reduced protein levels of RELA^I250V^ (Figure 3A). We postulated that reduced protein expression may be due to decreased stability of RELA^I250V^, however culturing with protease inhibitor MG132 did not rescue expression (Supplementary Figure 1C) suggesting an alternate mechanism of destabilisation. We also observed that RELA^D409N^ had a reduced molecular weight compared to RELA^WT^ (Figure 3A), potentially from differential post-translational modification. To rule out mislocalisation of mutant RELA, we expressed RELA^I250V^, RELA^R295H^ and RELA^D409N^ in HEK293 cells and stained using FLAG antibody. There was no obvious difference in localisation of the RELA variants compared with WT (Supplementary Figure 1D). To examine whether RELA protein expression was altered in patients, we immunoblotted lysates from patient PBMCs (Figure 3B). In contrast to the overexpressed alleles, there was no change in the expression of RELA in patient A.I.1 who is heterozygous for *RELA*^*I250V*^. Patient D.I.1 heterozygous for *RELA*^*P321A*^ appeared to have a slight reduction in RELA expression.

**Figure 3.**
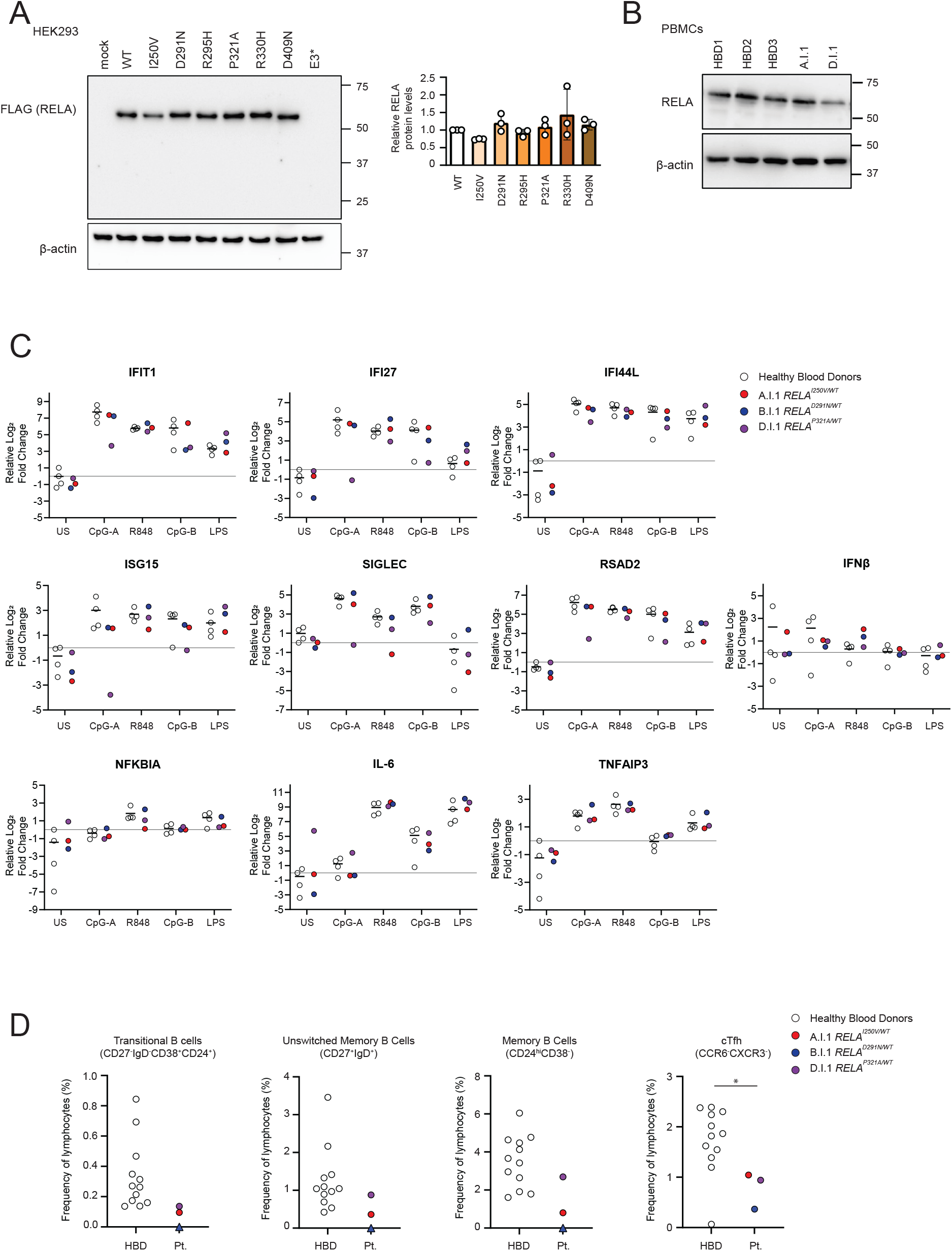
RELA expression and functional analysis in patients. (A) Immunoblot of RELA from transfected HEK293 cell lysates. Graph represents band densities relative to WT lane calculated from 3 independent experiments. (B) Immunoblot of endogenous RELA in PBMC lysates. (C) Relative expression of ISGs and NF-κB regulated genes in PBMCs from patients and healthy donors at baseline and after stimulation with TLR agonists R848 (TLR7/8), CpG-A and CpG-B (TLR9) and LPS (TLR4). Relative abundance of each transcript was normalized to *SDHA* transcripts and fold-change determined relative to a single calibrator. Median change displayed for healthy donors. (D) Immunophenotyping of patient PBMCs (triangle = too few events within gate, designated as 0%).

### Patients display a reduction in B and T cell subsets

Immunophenotyping was performed for patients A.I.1, B.I.2 and D.I.1 (Figure 3D). A reduction in the following B cells subsets was noted for patients A.I.1 and D.I.1 and for patient B.I.2 the populations were designated zero as too few events fell within the applied gates to confidently call population percentages; transitional B cells (CD27^-^ IgD^-^CD38^+^CD24^+^), unswitched memory B cells (CD27^+^IgD^+^), memory B cells (CD24^hi^CD38^-^). The percentage of the cTfh population defined as CCR6^-^CXCR3^-^ was significantly reduced for all patients.

### Patients do not display an elevated interferon stimulated gene (ISG) transcript signature

Leukocytes from patients with dominant negative *RELA* variants displayed type I interferon signatures and excessive type I/III interferon production following TLR7 stimulation ^6^. To determine if patients A.I.1, B.I.2 and D.I.1 demonstrated a similar bias toward IFN signalling we measured the induction of interferon stimulated genes and NF-κB responsive genes (Figure 3C) in PBMCs following stimulation with agonists for TLR7/8 (R848), TLR9 (CpG-A and CpG-B) and TLR4 (LPS). In unstimulated conditions, ISG transcripts were similar between patients and healthy blood donors, except for patient A.I.1 (*RELA*^*I250V/+*^) who displayed an elevated level of *IFNβ* transcripts compared to most healthy blood donors. D.I.1 had elevated *IL-6* transcript levels at baseline, however levels were comparable to healthy donors after NF-κB activation via R848, CpG-B and LPS. Following stimulation with TLR9 agonists CpG-A and CpG-B, D.I.1 displayed reduced induction of *IFIT1, IFI27, SIGLEC, RSAD2* and *ISG15* transcripts. The induction of ISGs and NF-κB responsive genes was similar between patients and controls following TLR7/8 stimulation with R848, except for A.I.1 who did not induce *SIGLEC* transcripts and had increased levels of *IFNβ* mRNA. In summary, although high baseline and TLR7/8-derived *IFNβ* transcripts were detected in A.I.1, an exaggerated ISG signature was not evident in patients following TLR activation *ex vivo*.

## Discussion

We have identified seven rare and/or novel *RELA* variants in patients with a range of immune diseases. To date, *RELA*^*E3\**^ is one of two reported *RELA* variants described in a patient diagnosed with autoimmune lymphoproliferative syndrome and is expected to result in *RELA* haploinsufficiency. Other manifestations present in the cohort investigated included lymphadenopathy, arthralgia, arthritis, and skin and gastrointestinal symptoms, which also align with phenotypes described in patients with *RELA* LoF and dominant negative mutations^6, 14^. Our data extends the *RELA*- associated clinical spectrum to include juvenile dermatomyositis, SLE and granulomatous features including a diagnosis of GPA and sarcoidosis.

Three *RELA* variants result in functional impairment of *NF-κB* activity, and our data suggests this occurs through different mechanisms. We were unable to detect RELA^E3*^ protein (data not shown) indicating that loss of activity is likely a direct result of RELA protein loss. Similarly, RELA^I250V^ expression was reduced, however we cannot discount that reduced activity stems from altered DNA binding. DNA binding and dimerisation is essential to downstream activation of NF-κB inflammatory genes^15^. RELA^I250V^ and RELA^R295H^ are both located in the DNA binding domain and both mutant proteins lost activity in homodimer and RELA/p50 heterodimer form. Additionally, PTMs regulate NF-κB DNA binding and protein-protein interactions ^15^. Our data reveal that D409N altered RELA molecular weight suggestive of an altered PTM. Further investigation is needed to determine whether protein-protein interactions are altered for RELA^D409N^.

In contrast to the hypomorphic activity towards the NF-κB response element, RELA^I250V^ and RELA^R295H^ (with the p300 coactivator) displayed elevated activity on the *IFNβ* promoter, and elevated basal *IFNβ* transcripts were detected in PBMCs from the patient heterozygous for RELA^I250V^. RELA is an important transactivator as a component of the IFNB enhanceosome^11^ and IFN gene induction requires timely recruitment of RELA ^16^. Others have demonstrated that murine RelA potently synergises with Irf3 to promote the induction of the *Ifnβ* promoter^15^. Here we observed the opposite, demonstrating that the 448 amino acid RELA^WT^ isoform which lacks IRF binding sites suppresses IRF-3, -5 and -7 induced *IFNβ* promoter activity. This reveals that there may be isoform-specific regulatory functions. We demonstrate that RELA^I250V^ and RELA^R295H^ proteins have reduced capacity to suppress IRF activity. The mechanism driving the loss of RELA suppressive activity is unknown but may occur via changes in non-canonical NF-κB signalling. The non-canonical NF-κB signalling pathway negatively regulates *Ifnβ* induction by suppressing histone modification and attenuates RelA recruitment to the *Ifnβ* promoter, likely because of competitive binding with the non-canonical RelB/p52 dimer^15^.

In conclusion, we identify new rare and novel *RELA* variants across a broader spectrum of clinical syndromes. Whilst these *RELA* variants have common pathophysiological markers of NF-κB LoF, they have distinct impacts of IFN functionality. These findings highlight the important role of RELA in a wide range of autoimmune and autoinflammatory diseases.

## Acknowledgements

The authors would like to thank the staff of the Australian Cancer Research Foundation Biomolecular Resource Facility (ACRF-BRF, JCSMR, Australian National University) for Sanger sequencing. This work was supported by funding to SJ from the Medical Research Future Fund and NHMRC (MRF2025693 and MRF20223294), Canberra Hospital Private Practice Fund, Jacquot Research Establishment and Career Development Award, and the Hindmarsh Family and McCusker Charitable Foundations.

## Figure Legends

**Supplementary Figure 1** (A) NF-κB luciferase reporter activity in HEK293 cells transfected with 25 ng of *WT* or *MT RELA* plasmids and 50 ng of *p300*. Graph represents 2 independent experiments. (B) Immunoprecipitation of FLAG tagged RELA in HEK293 cells co-transfected with *WT* or *MT RELA* and *IRF7* plasmids. Membranes were immunoblotted using anti- FLAG, anti-IRF7 and anti-GAPDH antibodies. Input lysates were run on the same gel but were non-contiguous (black borders). (C) Transfected HEK293 cells expressing RELA^WT^ or RELA^I250V^ with and without MG132 treatment. Membranes were immunoblotted with anti-FLAG anti-pan ubiquitin and anti-βactin antibodies. The increase in ubiquitin with MG132 treatment supports proteasome inhibition. (D) Immunofluorescence staining of WT and MT RELA in transfected HEK293 cells.

## Notes

### Competing Interest Statement

The authors have declared no competing interest.

## References

1. Pisetsky DS. Pathogenesis of autoimmune disease. Nat Rev Nephrol 2023;19(8):509–524.

2. Zhang Q, Lenardo MJ, Baltimore D. 30 years of NF-κB: a blossoming of relevance to human pathobiology. Cell 2017;168(1):37–57.

3. Barnabei L, Laplantine E, Mbongo W, et al. NF-κB: At the Borders of Autoimmunity and Inflammation. Front Immunol 2021;12:716469.

4. Comrie WA, Faruqi AJ, Price S, et al. RELA haploinsufficiency in CD4 lymphoproliferative disease with autoimmune cytopenias. J Allergy Clin Immun 2018;141(4):1507-1510. e1508.

5. Lecerf K, Koboldt DC, Kuehn HS, et al. Case report and review of the literature: immune dysregulation in a large familial cohort due to a novel pathogenic RELA variant. Rheumatology 2023;62(1):347–359.

6. Moriya K, Nakano T, Honda Y, et al. Human RELA dominant-negative mutations underlie type I interferonopathy with autoinflammation and autoimmunity. J Exp Med 2023;220(9):e20212276.

7. Hayden MS, Ghosh S. Shared principles in NF-kappaB signaling. Cell 2008;132(3):344–362.

8. Field MA, Cho V, Andrews TD, et al. Reliably detecting clinically important variants requires both combined variant calls and optimized filtering strategies. PloS one 2015;10(11):e0143199.

9. Gibson G. Rare and common variants: twenty arguments. Nature Reviews Genetics 2012;13(2):135–145.

10. Sun S-C. The non-canonical NF-κB pathway in immunity and inflammation. Nat Rev Immunol 2017;17(9):545–558.

11. Panne D, Maniatis T, Harrison SC. An atomic model of the interferon-β enhanceosome. Cell 2007;129(6):1111–1123.

12. Pascual V, Farkas L, Banchereau J. Systemic lupus erythematosus: all roads lead to type I interferons. Curr Opin Immunol 2006;18(6):676–682.

13. Fan S, Popli S, Chakravarty S, et al. Non-transcriptional IRF7 interacts with NF-κB to inhibit viral inflammation. J Biol Chem 2024;300(4).

14. An JW, Pimpale-Chavan P, Stone DL, et al. Case report: Novel variants in RELA associated with familial Behcet’s-like disease. Front Immunol 2023;14:1127085.

15. Jin J, Hu H, Li HS, et al. Noncanonical NF-κB pathway controls the production of type I interferons in antiviral innate immunity. Immunity 2014;40(3):342–354.

16. Wang J, Basagoudanavar SH, Wang X, et al. NF-κB RelA subunit is crucial for early IFN-β expression and resistance to RNA virus replication. J Immunol 2010;185(3):1720–1729.

