## Supplementary Tables 1-3, Figure 1, Methods for "Rare and novel *RELA* variants are common in systemic autoimmunity"

Supplementary Table 1. Patient demographics and clinical history

| **Patient** | **AA change** | **AA position** | **Sex** | **Age (years)** | **Clinical history** |
|---|---|---|---|---|---|
| A.I.1 | I>V | 250 | Female | 36 | SLE |
| B.I.1 | D>N | 291 | Male | 79 | Sarcoidosis |
| C.II.1 | R>H | 295 | Female | 44 | Fibromyalgia, GPA, Seronegative RA, CVID - hypogammaglobulinaemia. |
| *Father (C.I.1) has IgG subclass deficiency and history of childhood bronchiectasis, lobectomy, sinus surgery due to recurrent sinusitis (opportunistic fungal infection detected). | | | | | |
| D.I.1 | P>A | 321 | Female | 42 | Scleroderma, Spondyloarthritis |
| E.II.1 | R>H | 330 | Female | - | SLE glomerulonephritis syndrome |
| F.II.2 | D>N | 512 | Male | 26 | Juvenile dermatomyositis |
| G.II.1 | E>Ter | 3 | Female | 13 | ALPS |

AA=Amino Acid; SLE = Systemic lupus erythematosus; GPA = Granulomatosis with polyangiitis; RA = Rheumatoid arthritis; CVID= Common Variable Immunodeficiency CENP = Centromere protein; ALPS = Autoimmune lymphoproliferative syndrome.

Supplementary Table 2. Single nucleotide variants and pathogenicity predictions

| Nucleotide change/ Genomic Coordinates/ rs identifier | Mutation type | Minor allele frequency (gnomAD) | PolyPhen Score | SIFT score | CADD (Phred) score |
|---|---|---|---|---|---|
| c.749 A>G Chr11:65658416 | Missense | Novel | 0.71 | 0.02 | 24.5 |
| c.872 G>A Chr11:65658293 rs61759893 | Missense | Rare MAF: 0.004278 | 0.54 | 0.01 | 24.2 |
| c.884 G>A Chr11:65655929 rs150601443 | Missense | RARE MAF: 0.000261 | 0.54 | 0.18 | 23.9 |
| c.962 C>G Chr11:65655760 | Missense | RARE MAF: 0.000001239 | - | - | 21.8 |
| c.998 G>A Chr11:65655732 rs1232341468 | Missense | RARE MAF: 0.000007435 | 0.66 | 0.03 | 26.5 |
| c.1226 G>A Chr11:65654500 rs1463059248 | Missense | RARE MAF: 0.000004997 | 0.73 | 0.03 | 25 |
| c.8 G>T Chr11:65662826 rs1856608678 | Nonsense | NOVEL | - | - | 55 |

Supplementary Table 3. Predicted disruption to RELA structure and stability

| ***RELA* variant** | **DynaMut (ddG)** | **Grantham's Distance Score** | **Located in a disordered region (yes/no)** | **Localisation: Surface or buried (S/B)** | **AlphaFold pLDDT Score** |
|---|---|---|---|---|---|
| I250V | -0.939 | 29 | no | B | 98.70 |
| D291N | 0.018 | 23 | no | S | 86.09 |
| R295H | 0.184 | 29 | no | S | 72.71 |
| P321A | 0.233 | 27 | yes | S | 53.54 |
| R330H | 0.384 | 29 | yes | S | 54.07 |
| D512N | 0.167 | 23 | yes | S | 45.26 |

ddG=delta delta G; pLDDT=predicted local distance difference test

### Supplementary Figure 1

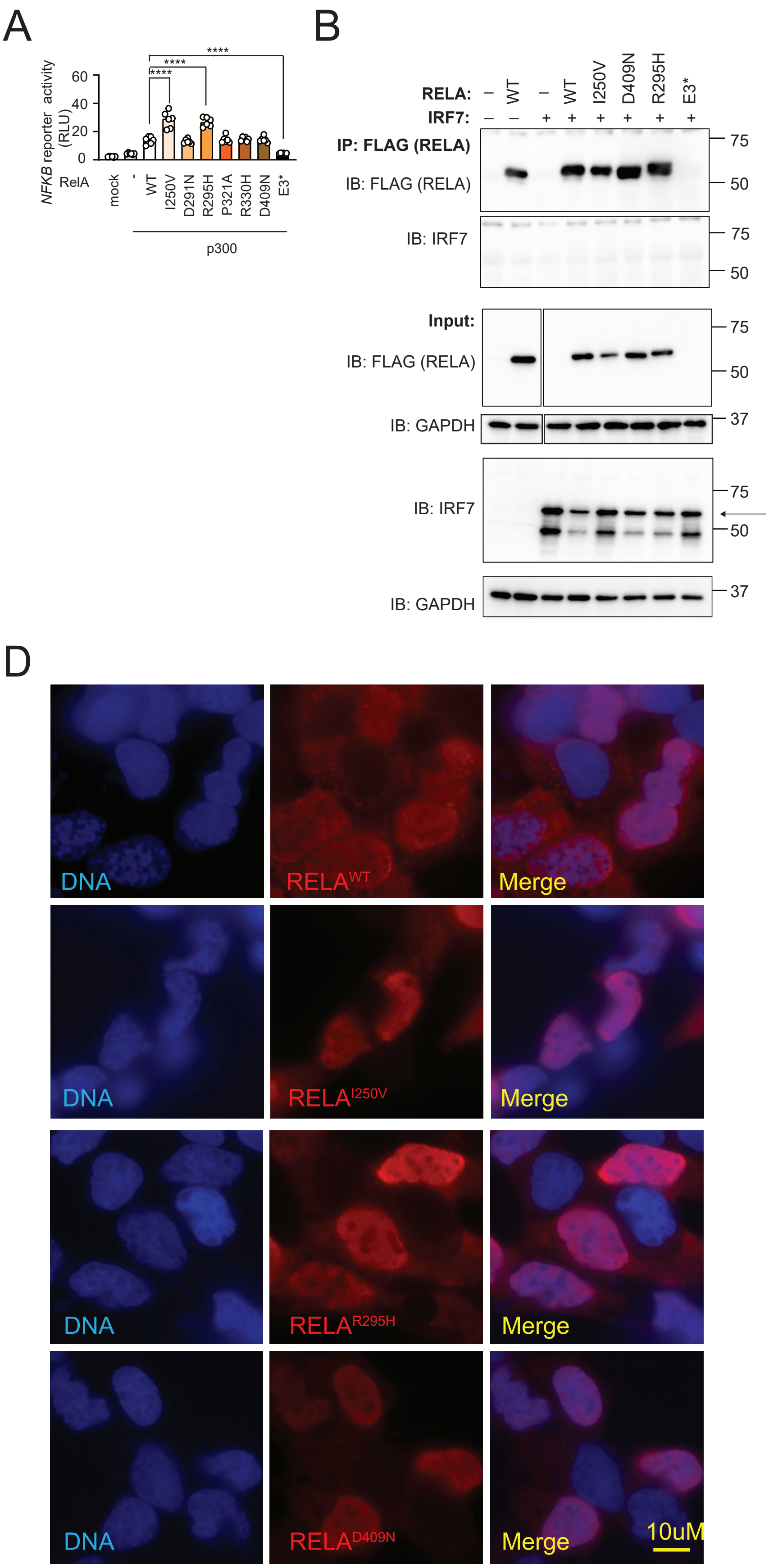

### Supplementary Methods

#### Sanger sequencing

Genomic DNA was isolated from frozen PBMCs using the QIAamp DNA Blood Mini Kit (QIAGEN) following the manufacturer's instructions. DNA regions containing variants of interest were amplified via PCR using Q5 Hot Start High-Fidelity Master Mix (New England Biolabs) according to manufacturer's instructions. PCR products were isolated via agarose gel electrophoresis and purified using the Wizard® SV Gel and PCR Clean-Up System (Promega). Sanger sequencing was performed by the Biomolecular Resource Facility, ANU. Sequencing products were analysed on a 3730 DNA Analyser (Applied Biosystems) and variants analysed using SnapGene (Version 7.0.3, GSL Biotech LLC).

#### Expression plasmids and mutagenesis

The *WT RELA* coding sequence was subcloned from pDONR223_RELA_WT (gifted by Jesse Boehm & William Hahn & David Root, Addgene #82160) into the destination mammalian expression vector pDEST-FLAG (gifted by Emanuele Panza, Addgene #183511) using the Gateway LR Clonase II system (Invitrogen). *MT RELA* plasmids were created using the Q5® Site-Directed Mutagenesis Kit (New England Biolabs) according to the manufacturer's instructions. Other plasmids used include p50 (gifted by Warner Greene, Addgene #21965), IRF3 (SinoBiological, HG12007-UT), IRF5 (OriGene Technologies #SC104269), MyD88 (GenScript #OHu21475), p300 (gifted by Tso-Pang Yao, Addgene #30489 and pcDNA3.1+ (Invitrogen). The open reading frame from IRF7 plasmid (Origene Technologies #SC125442) was subcloned into pcDNA3.1+ (Invitrogen) between restriction sites *HindIII* and *MfeI* using T4 DNA ligase (Promega).

**Antibodies**

The following antibodies were used for immunoblotting, co-immunoprecipitation and immunofluorescence assays: mouse anti-FLAG® M2 (#F1804, Sigma), rabbit anti-NF-κB p65 (D14E12) (Cell Signalling Technology), rabbit anti-ubiquitin (E4I2J) (#43124, Cell Signalling Technology), rabbit anti-IRF7 (G-8) (#sc-74472,Santa Cruz), mouse anti-GAPDH (6C5) (#8245, Abcam), rabbit anti-β-Actin (13E5) (#5125, Cell Signalling Technology) and Alexa Fluor 568 donkey anti-mouse IgG (#A10037, Invitrogen).

**Quantitative PCR (qPCR)**

PBMCs were stimulated with LPS (lipopolysaccharide) (Sigma), Resiquimod (R848) (Invivogen), CpG-A (ODN-2216) or CpG-B (ODN-2006) (Miltenyi Biotec) for 24 hours. RNA was extracted from PBMCs using Trizol (Sigma). cDNA was synthesized using SuperScript IV Reverse Transcriptase (Thermo Fisher Scientific). qPCR was performed with the QuantStudio™ 12K Flex System (Thermo Fisher Scientific) using the SYBR green method following manufacturer's instructions. Primer sequences are listed in Supplementary Table 1. Relative expression was calculated using the 2−ΔΔCt method.

List of qPCR primers used:

| TARGETS | | PRIMERS |
|---|---|---|
| SIGLEC | Forward | 5'TGGAGAAGGAGGCGTGTTTG3' |
| | Reverse | 5'AGGATCAATGAGCTGCGTGG3' |
| ISG15 | Forward | 5'ACAGCCATGGGCTGGGA3' |
| | Reverse | 5'CCTTCAGCTCTGACACCGAC3' |
| IFIT1 | Forward | 5'ATTTACAGCAACCATGAGTACAAA3' |
| | Reverse | 5'TCCCACACTGTATTTGGTGTCT3' |
| IFNB | Forward | 5’ GACCATCTATGAGATGCTCCAGAACA3’ |
| | Reverse | 5’ CAGGAGGTTCTCAACAATAGTCTCAT3’ |
| IFI44L | Forward | 5'GACTTCTCAAAGCCGGGTCA3' |
| | Reverse | 5'CCTTCATGGGGTCCAGTTCC3' |
| RSAD2 | Forward | 5'CCTGTCCGCTGGAAAGTGTT3' |
| | Reverse | 5'GACACTTCTTTGTGGCGCTC3' |
| IL-6 | Forward | 5'AGAGGCACTGGCAGAAAACA3' |
| | Reverse | 5'TCACCAGGCAAGTCTCCTCA3' |
| NFKBIA | Forward | 5'CCCTACACCTTGCCTGTGAG3' |
| | Reverse | 5'TAGACACGTGTGGCCATTGT 3' |
| TNFAIP3 | Forward | 5'CTGAAAACGAACGGTGACGG 3' |
| | Reverse | 5'GCAAAGCCCCGTTTCAACAA 3' |
| IFI27 | Forward | 5'CTCCTTCTTTGGGTCTGGCT 3' |
| | Reverse | 5'GGCCACAACTCCTCCAATCA 3' |
| SDHA | Forward | 5'TGGCATTTCTACGACACCGTG3' |
| | Reverse | 5'GCCTGCTCCGTCATGTAGTG3' |

**Co-immunoprecipitation**

HEK293 cell extracts were prepared using 1% Triton X-100 (Merck) lysis buffer (25 mM Tris-HCl, 150 mM sodium chloride, 1 mM EDTA, 5% glycerol), containing PhosSTOP (Roche) and cOmplete protease inhibitor (Roche). FLAG-RELA proteins were immunoprecipitated using Anti-FLAG® M2 Affinity Gel (Millipore). Immunoprecipitated proteins and WCE were separated via denaturing SDS-polyacrylamide gel electrophoresis and transferred to nitrocellulose membranes. Membranes were probed for FLAG-RELA and IRF7 using appropriate primary and secondary antibodies. Membranes were developed with Clarity Western ECL

Substrate (Bio-Rad Laboratories) and imaged on ChemiDoc System (Bio-Rad Laboratories).

**Immunofluorescence staining and microscopy**

HEK293 cells were seeded onto coverslips and transfected with the relevant plasmids. Post transfection, cells were fixed in 3.7% formaldehyde, permeabilized with 1% Triton X-100, and blocked with 5% bovine serum albumin (BSA)/0.1% Triton X-100 for 1 hour. Cells were stained with indicated primary antibodies in blocking buffer overnight, and secondary Alexa Fluor 568 donkey anti-mouse IgG before mounting in VectaShield Antifade mounting medium with DAPI (Vector Labs). Images were captured on an Olympus IX71 inverted microscope with 100X oil objective PlanApo N (1.40 NA/ 0.17 WD) using DP Controller software and compiled using Adobe Photoshop.
